# Advancing quality-control for NGS measurement of actionable mutations in circulating tumor DNA

**DOI:** 10.1101/2021.04.06.438497

**Authors:** James C. Willey, Tom Morrison, Brad Austermiller, Erin L. Crawford, Daniel J. Craig, Thomas M. Blomquist, Wendell D. Jones, Aminah Wali, Jennifer S. Lococo, Nathan Haseley, Todd A. Richmond, Natalia Novoradovskaya, Rebecca Kusko, Guangchun Chen, Quan-Zhen Li, Don Johann, Ira W. Deveson, Tim Mercer, Leihong Wu, Joshua Xu

## Abstract

The primary objective of the FDA-led Sequencing and Quality Control Phase 2 (SEQC2) project is to develop standard analysis protocols and quality control metrics for use in DNA testing to enhance scientific research and precision medicine. This study reports a targeted next generation sequencing (NGS) method that enables more accurate detection of actionable mutations in circulating tumor DNA (ctDNA) clinical specimens. This advancement was enabled by designing a synthetic internal standard spike-in for each actionable mutation target, suitable for use in NGS following hybrid-capture enrichment and unique molecular index (UMI) or non-UMI library preparation. When mixed with contrived ctDNA reference samples, internal standards enabled calculation of technical error rate, limit of blank, and limit of detection for each variant at each nucleotide position, in each sample. True positive mutations with variant allele fraction too low for detection by current practice were detected with this method, thereby increasing sensitivity.

## INTRODUCTION

Measurement of circulating tumor DNA (ctDNA) in blood samples with recently developed next generation sequencing (NGS) provides a non-invasive “liquid biopsy” to detect and genetically target cancer as well as to monitor response to treatment^1–10^. The U.S Food and Drug Administration (FDA) recently approved certain ctDNA assays for diagnostic applications. However, the approved assays are limited by current common practice. For example, a limit of detection (LOD) is estimated *a priori* for each type of actionable mutation, and then a constant predefined value is applied to every sample. For a commercial platform recently approved by the FDA, the predefined variant allele fraction (VAF) LOD for measurement of an actionable BRAF mutation was 1.1% or 0.2% for DNA input of 5 ng or 30 ng respectively^11^. However, there is both variant-specific and inter-sample variation in LOD^12, 13^. Thus, application of a predefined VAF LOD to analysis of every actionable mutation in every sample is limiting because variants with lower VAFs could be reliably measured in many ctDNA samples. Further, even “conservative” predetermined LOD thresholds may not ensure reproducible results due to insufficient measurement of, and control for, sample-specific and site-specific technical error^12, 13^ and/or stochastic sampling associated with small and/or poor quality ctDNA samples^9, 13–19^. In order to obtain maximum reliable information from limited ctDNA samples, there is a need for methods that provide sample- and variant-specific LOD, confidence limits for each test result, and inter-laboratory concordance^13, 16, 20–22^. The method described here was designed to enable sample- and variant-specific LOD while ensuring reliable reporting of NGS VAF data. This study was conducted as a collaborative inter-laboratory effort through the FDA-led Sequencing Quality Control Consortium Phase 2 (SEQC2) project^23^.

Most false positive variants are caused by NGS technical errors due to regional and site-specific chemical and physical factors during library preparation and sequencing^12,13^. Multiple approaches have been developed to address this^24–29^. One approach is to capture recurrent technical artifacts by sequencing many normal samples, then apply a variant-specific LOD value established *a priori* based on the rate of technical error that gives rise to each respective variant^25^. This approach typically requires sequencing data from more than 40 normal samples and is better suited to highly automated reference laboratories with sufficient testing bandwidth to regularly confirm that the technical error has not altered due to small drifts in reagents, instruments, and operators. Further, this approach does not control for inter-measurement variation in sample-specific inhibitors, instrument performance, or operator proficiency. Another approach is to reduce sequencing artifacts by attachment of a unique molecular identifier (UMI) to each DNA molecule during library preparation, followed by bioinformatic identification of consensus among PCR replicates^24, 26–28^. This method facilitates identification and removal of technical errors that are inconsistent with the consensus, prior to variant calling. However, UMI methods may introduce new non-systematic errors, unrelated to chemical and physical technical errors, that must be identified and filtered empirically^29^. It is reasonable to hypothesize that the combination of known chemical-physical sources and potential non-systematic bioinformatic sources of technical error define the limit of the blank (LOB) for measurement of each variant. Further, it is reasonable to hypothesize that the primary determinants of LOD and confidence limits in each measurement of a variant are a) variant-specific LOB, and b) the number of ctDNA sample variant copies captured in the library preparation.

We previously demonstrated that Standardized Nucleic Acid Quantification (SNAQ-SEQ) internal standard (IS) controls improved accuracy of variant calling with amplicon-based sequencing^12, 13^. Specifically, systematic chemical-physical technical errors were closely modeled by synthetic IS spike-in molecules added to each sample^12,13^. Because IS controls were known to be reference genome sequence, any variant detected in IS reads was caused by technical errors accrued in the NGS testing process and the technical error fraction for each detected IS variant closely reflected the error fraction for that same variant in the native template sequence of the sample tested. As such, when IS was added to a sample, the variant-specific technical error in IS provided a basis for establishment of variant-specific LOD in that sample.

There is an unmet need for hybrid capture targeted NGS methods that provide an LOD for measurement of each actionable mutation in each ctDNA sample. To address this need, we implemented a new design for SNAQ-SEQ IS; a design suitable for use in NGS following hybrid-capture enrichment and unique molecular index (UMI) or non-UMI library preparation. In assessment of SNAQ-SEQ IS clinical utility, targeted NGS libraries were prepared with UMI following hybrid enrichment because this is the predominant NGS method used by the majority of oncopanels developed for ctDNA analysis^30^.

The purpose of this study was to a) evaluate whether synthetic spike-in IS control for technical errors associated with hybrid capture, b) to assess utility of synthetic spike-in IS as an orthogonal quality-control for UMI library analysis, and c) to determine whether mixture of IS with each sample interferes with existing hybrid capture workflow or cause additional errors. It was necessary to develop bioinformatic methods that efficiently separate the re-designed spike-in IS from sample native template (NT) sequence reads in FASTQ files prior to pipeline analysis and variant calling. Moreover, this approach required development of a statistical approach for variant calling based on comparative analysis of sequencing reads from each sample DNA NT and respective IS. In order to test the clinical utility of methods developed through these studies, we also prepared contrived ctDNA reference samples ^30^ with ground truth true positive (TP) variant sites and ground truth true negative (TN) reference sites. The goal was to determine whether this method would improve quality control in NGS measurement of actionable mutations in ctDNA specimens and thereby enable increased clinical sensitivity without loss of specificity.

## RESULTS

### A ground truth dataset was established in SEQC2 reference *Sample A*

The ground truth data set created in a related SEQC2 project study^30, 31^ was extended in this study to a 6.8 kb region common to the SNAQ-SEQ IS mixture (described in **Methods**), the Illumina TST170 panel, and Roche SeqCap EZ Choice custom PHC panel. The Roche SeqCap EZ Choice custom PHC panel was used for hybrid capture enrichment followed by non-UMI library preparation. Briefly, ground truth TP variants were established by sequencing samples from a dilution series in which an SEQC2 contrived tumor sample (SEQC2 reference *Sample A*) was diluted with a normal sample (SEQC2 reference *Sample B). Sample A* was a mixture of 10 cancer cell lines. TP were identified as variants that responded appropriately to titration. Analysis of the *Sample-A/B* serial dilution samples expanded the set of TP in the 6.8 kb consensus region to 28, an increase from 8 reported in a related SEQC2 study^30^ **(Supplemental Figure 1)**. Of the 6.8 kb region covered, 3500 positions had >1000 coverage in the ground truth data, sufficient to establish them as TN. As such, any suspicious variant call in a test sample at one of these 3500 positions could be confirmed as false positive (FP) if there was not a TP variant at the same position in the ground truth data (i.e., demonstrated lack of titration response).

### Variant calling by SNAQ-SEQ QC with Poisson Exact Test (PET) analysis

As detailed in the **Methods** section, SEQC2 Reference *Sample A* was mixed with SNAQ-SEQ IS mixture then subjected to hybrid capture target enrichment with the Roche SeqCap EZ Choice custom PHC panel, non-UMI library preparation, and sequencing. As presented in **Figure 1a**, a FASTQ file was bioinformatically separated into IS and NT bins through alignment to respective IS-specific and NT-specific Hg19 reference genomes. Precision for separation of NT from IS reads was >99.9997% (i.e., <0.0003% IS base changes in NT reads. Next, the respective IS FASTQ and NT FASTQ files were mapped with BWA MEM and positionally deduplicated with Picard Mark Duplicates. As an example of coverage covariation for IS and NT DNA fragments during enrichment, the IS and NT sequences mapped to *EGFR* gene are graphed in **Figure 1b**. The histogram demonstrates that, although capture efficiency and subsequent sequence coverage varied considerably for each probe region, the IS (identified by presence of synthetic dinucleotide variants used to separate IS from NT) and NT were captured proportionately across all probes in the *EGFR* panel with peak capture efficiency in exonic regions.

**Figure 1.**
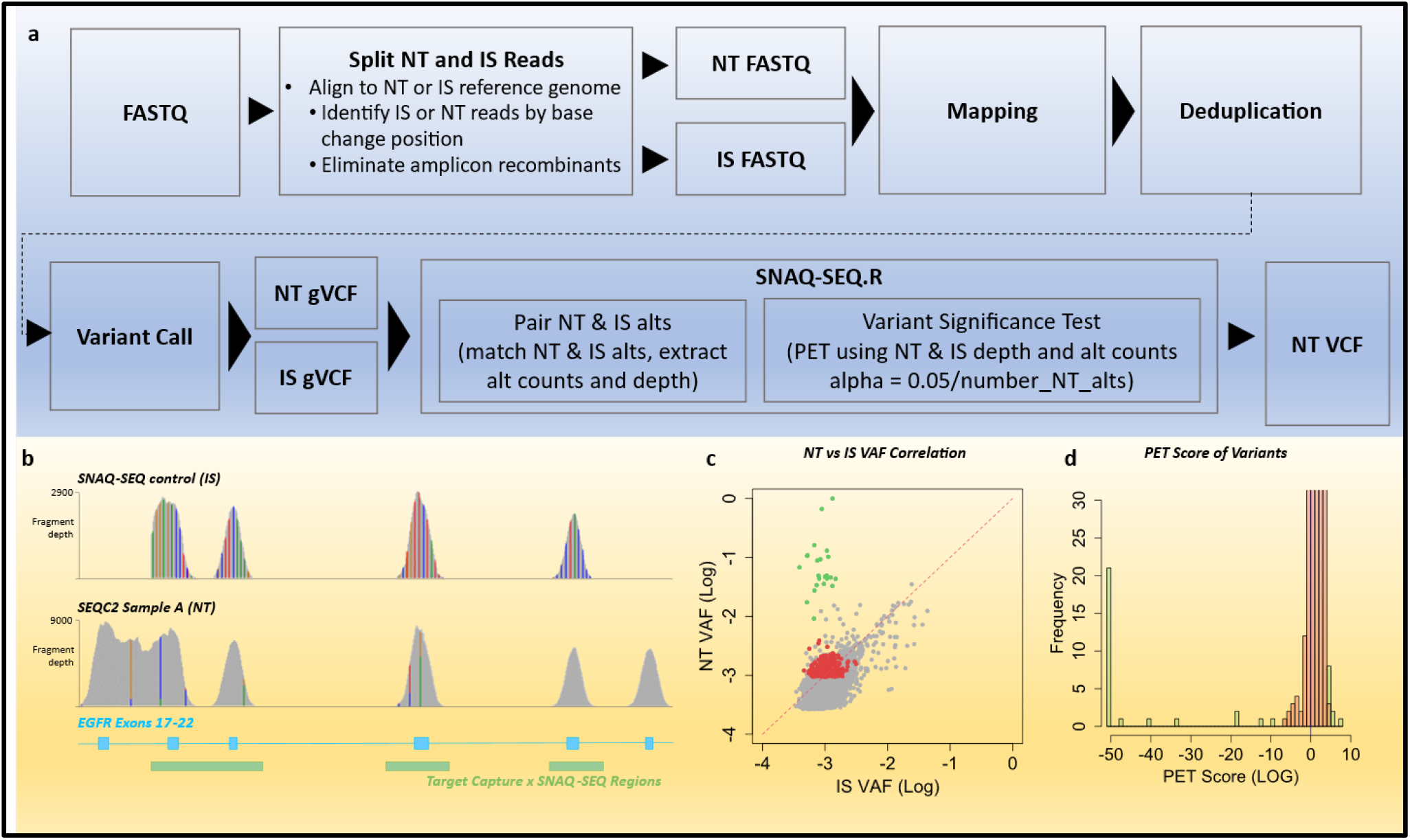
SNAQ-SEQ Method. **a**. Integration of SNAQ-SEQ into NGS pipeline (see details in Methods). **b**. Analysis of *EGFR* region in SEQC2 reference *Sample A* mixed with SNAQ-SEQ IS followed by enrichment on the Roche SeqCap EZ Choice custom PHC panel. Histogram of *EGFR* region (x-axis) vs fragment depth (coverage) (y-axis) for internal standard (IS) (top) and native template (NT) (bottom) sequence reads. Colored columns indicate bases alternate (alt) to hg19 reference with >5% VAF. IS alts were engineered dinucleotides, NT alts were SNPs derived from one or more cancer cell lines in *Sample A*. The three sequence regions spanned by IS (green bars) overlap 4 of the 6 depicted *EGFR* exons (blue bars) bounded by the Roche panel target regions. **c**. The 5608 Sample A NT alts with matched alts in IS sequence were plotted: NT VAF (y-axis) and IS VAF (x-axis). NT alts called by VarDict are indicated as significant (green filled) or not significant (red filled) by SNAQ-SEQ QC PET analysis, and alts that were not called by VarDict or SNAQ-SEQ QC PET analysis are indicated as grey circles. Red dashed line is 1:1 correlation in VAF value. **d**. Frequency (y-axis) of alts binned according to PET score (x-axis) for the 374 NT alts called by VarDict (left side of plot, scores <0) and 895 IS alts (right side of plot, scores >0), with significant alts as green columns, non-significant as red columns.

VarDict was used to call each IS and NT variant (or alternative [alt]) allele using the IS and NT reference sequences, respectively, then the pileup option was used to create IS and NT genome (g)VCF files. SEQC2 *Sample A* libraries yielded 16,310 NT alt bases at 8,571 positions within exons spanned by IS (other than IS dinucleotide sites). Of these *Sample A* NT alts, 5608 had the same alt in the IS sequence and this allowed bivariate plot of NT VAF (y-axis) and IS VAF (x-axis) (**Figure 1c)**. Among the 16,310 NT alt bases, VarDict called 374 alts (i.e. PASS FILTER). Other than the expected 28 TP variants, the remaining VarDict PASS variants (N = 346) arose among the 3,500 TN positions and, by definition, were FP arising from technical errors.

SNAQ-SEQ QC used both the VarDict IS gVCF and NT gVCF files to determine significance of each variant in the NT gVCF file. PET was used to analyze the relative VAF values for each NT alt and corresponding IS alt to establish LOB and LOD for measurement of the NT alt. Specifically, for each NT alt allele called by VarDict, statistical significance was determined by PET analysis of the number of alt observations (i.e., alt count and position coverage) for the respective NT and IS sequences. PET was an appropriate statistical test for this application because Poisson distribution best represents both a) the low number of NT alts commonly observed in small and/or degraded DNA samples, such as ctDNA and cytologic specimens, and b) the low number of NT and IS alts resulting from technical error^12^. Frequencies of the 374 VarDict PASS NT alts, binned according to PET score are presented in **Figure 1d**. VarDict **NT** PASS alts had PET scores less than zero (left side of plot) and VarDict **IS** PASS alts had PET scores greater than zero (right side of plot). Alts with PET score significantly different from NGS background error appear as green bars and those not distinguishable from NGS background error appear as red bars. Because each IS was synthesized and sequence verified, any IS alts, including those with significant PET score (green bars) relative to NT background, were technical error (FP). Therefore, the highest PET score observed for an IS technical error variant was used to determine PET score significance threshold for NT variants.

The 28 ground truth TP variants were called by VarDict, confirmed by SNAQ-SEQ QC with PET analysis (green columns with negative PET score in **Figure 1d**), and appear as green circles in **Figure 1c**. In contrast, 346 alts called by VarDict (VarDict FP) were correctly classified as TN by SNAQ-SEQ QC (red columns in **Figure 1d**, and red circles in **Figure 1c**). The remaining TN alts were correctly not called by VarDict or SNAQ-SEQ QC analysis and appear as grey circles in **Figure 1c**. As is evident in **Figure 1c**, genome positions with matched alts in NT and IS, and NT alt confirmed as TN by SNAQ-SEQ QC (grey and red circles) clustered around the 1:1 correlation line. Further, variation in NT and IS VAF among these technical error variants ranged over three logs (base10) consistent with amplicon library data in prior reports^12, 13^. This high correlation between SNAQ-SEQ IS sequences and corresponding human DNA sample NT sequences at TN sites enabled use of IS as an internal negative control to estimate NGS background.

### Evidence consistent with no effect of SNAQ-SEQ IS on vendor variant calling performance

The 6.8 kb consensus region spanned by SNAQ-SEQ IS represented <4% of the 150 to 500 kb regions spanned by hybrid capture panels used in this study and therefore comprised a small fraction of total library reads. Further, once the IS reads were separated from NT, the presence of the controls during library preparation had no measurable effect on reproducibility, sensitivity, or false positive rate^30^. Based on these criteria, inclusion of SNAQ-SEQ IS with SEQC2 contrived ctDNA reference samples did not interfere with hybrid capture library preparation platforms used in the SEQC2 study^30^.

### Demonstration of SNAQ-SEQ QC Utility: SNAQ-SEQ QC improved variant calling in SEQC2 contrived reference ctDNA

The SEQC2 contrived ctDNA reference samples used in this study closely resemble those used in a related SEQC2 study^30^ **(Methods** section). *Sample A* was a mixture of 10 cancer cell lines. *Samples D* and *E* were created through 5-fold or 25-fold dilution of *Sample A* DNA with normal cell line *Sample B* DNA. Therefore, *Samples D* and *E* have the same variants as *Samples A* and *B* but at different variant allele frequencies (see details in **Methods**)^30, 31^. As described in the related manuscript, SEQC2 *Samples B, D* and *E* were subjected to enzymatic fragmentation followed by size selection to create *Samples Bf, Df* and *Ef*^30^. The samples named *Df* and *Ef* in this study were derived from the same batch of material as the SEQC2 reference ctDNA samples (*ct-high* and *ct-low*) respectively, used in a related study from the SEQC2 project and VAF values were expected to be equivilent^30^.

*Samples Bf, Df*, and *Ef* were each mixed with aliquots of the fragmented IS spike-in mixture to make *Samples BfIS, DfIS* and *EfIS*, respectively (see **Methods**). The fragment size distribution of SNAQ-SEQ IS controls closely paralleled that of the SEQC2 reference samples (**Supplemental Figure 2**). Based on the ground truth data, following 5-fold dilution of *Sample A* in *Sample B, Sample DfIS* was expected to have 20 TP variants ranging from 0.5% to 1% VAF. Assuming good TST170 library preparation capture efficiency, the four *Sample DfIS* replicates at 50 ng input each (14,500 haploid genomes) would provide sufficient read counts for detection of each variant in the 0.5% to 1% range. In contrast, in *Sample EfIS* these same variants would range from below 0.1% to 0.5% VAF and those at the lower VAF range would challenge the detection limit.

Based on analysis of *Sample A*, an IS:NT ratio of at least 2.5:1 enabled better PET analysis due to better estimate the NGS error rate. Thus, the SNAQ-SEQ IS mixture was mixed with *Samples DfIS* and *EfIS* to achieve at least a 2.5:1 IS:NT ratio. The TST170 UMI pipeline called variants with a UMI-deduplicated read count of at least three. Assuming no variant was detected in IS, a variant with three NT alts would have a significant PET value (p<0.05) when the IS depth was 2.5-fold more than NT (i.e., there were sufficient observations of zero alts in IS to reach confidence that signal in NT was above noise). As presented in **Supplemental Table 1**, among the sequencing files for four replicate libraries of *Samples BfIS, DfIS*, and *EfIS*, across each position in the targeted regions the average IS:NT ratio was 4.8 and only 3% of NT alts had <2.5-fold IS coverage relative to NT. Further, there was high inter-library and inter-sample correlation for the IS/NT ratio measured for each of the targeted regions (mean CV 7.6%), as was observed for analysis of EGFR in *Sample A*, presented in **Figure 1b**. Inter-target variation in average IS/NT ratio ranged from 2.9 to 7.3-fold. This approximately 2.5-fold range in IS/NT ratio among the targeted regions arose from variation in concentration among the IS when the IS mixture was created.

Illumina UMI deduplication and variant calling was conducted to create VCF files for each replicate library, test for the presence of each expected TP, and assign a FILTER value for each detected TP: PASS or LowSupport. The Illumina pipeline reported only PASS variants. However, for purposes of this study, SNAQ-SEQ QC with PET analysis was applied to each alt with at least 3 alt counts and with either a PASS or LowSupport FILTER value, but not blacklisted. As presented for analysis of TP in *Sample A* **(Figures 1c and 1d)**, SNAQ-SEQ QC with PET analysis was used to estimate the technical error and calculate the LOB at each nucleotide position in the IS sequence. Graphs that present the distribution of PET scores for each variant in each replicate library of *Samples DfIS* and *EfIS* are depicted in **Figure 2a**. As is evident, in *Sample DfIS* (upper panel) the PET score corresponding to the TP variants were highly significant (green bars) and well separated from the background alts. Importantly, due to effective UMI suppression of technical error, no PET significant IS variants were observed in of the Sample DfIS or EfIS replicate libraries, in contrast to presence of PET significant IS variants in the non-UMI *Sample A* libraries (**Figure 1d**). An alternative view of the difference between TP and NGS sequencing error is depicted in the bivariate plots, **Figures 2b, c**. In **Figure 2b**, the VAF for each NT alt (y-axis) with a matching IS alt (x-axis) was presented. For *Sample DfIS* there were two well separated clusters. On the one hand, the VAF for each TP variant (circles), was two to three logs higher than for the matching IS alt. In contrast, the cluster associated with the 1:1 line comprises LowSupport alts that were not validated by PET analysis and therefore are not separable from technical error (red-filled triangles). In **Figure 2c** the IS VAF value is replaced with the LOB value estimated based on PET analysis. As with **Figure 2b**, the *Sample DfIS* TP variants cluster significantly above the LOB. In **Figure 2c** some *Sample DfIS* NT alts appear on the plot that were not present on **Figure 2b** because using PET analysis a LOB can be calculated even for NT alts for which zero alts were observed in IS. In contrast to *Sample DfIS* (**Figures 2b, c** upper panels), for *Sample EfIS* (**Figures 2b, c** lower panels) separation between TP (circles) and NGS error (triangles) was reduced due to the five-fold greater dilution of each *Sample A* TP variant in *Sample B* (see **Methods**). Correspondingly, the distribution of PET values for TP variants in *Sample EfIS* (**Figure 2a** lower panel) was closer to that for technical error.

**Figure 2.**
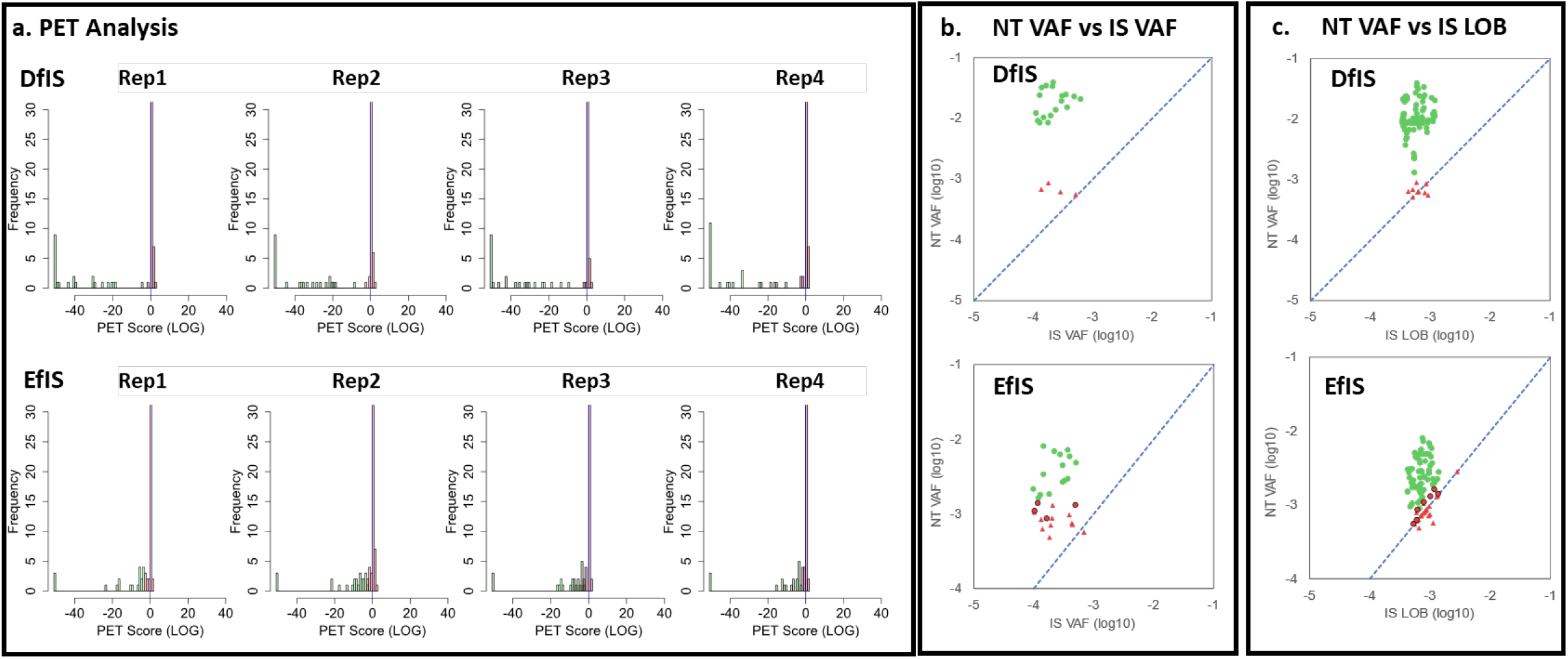
Effect of SNAQ-SEQ QC on ILM UMI VCF File Variants from *Samples DfIS* and *EfIS*. a. Poisson Exact Test (PET) analysis of ILM alts for each replicate library. PET score distribution of Illumina VCF file alts (alts with PMEAN >25 and PASS or LowSupport FILTER) for *Sample DfIS* (upper panels) and *Sample EfIS* (lower panels) replicate libraries; x-axis: PET score for NT alts (scores < 0) and IS alts (scores > 0); y-axis: alt frequency. SNAQ-SEQ QC PET analysis indicated alts significantly above background NGS technical error (green bars) or not significant (red bars). **b**. **Bivariate analysis of paired NT and IS alt VAF values**. NT alt VAF (y-axis) vs matched IS alt VAF in (x-axis) plotted for each NT alt. Circles: Ground true positive (TP) variants; green-filled PET significant and red-filled non-significant; Triangles: ILM FILTER LowSupport VCF that were not PET significant. The dashed reference line is the 1:1 correlation between NT and IS VAF values. **c. Bivariate analysis of paired NT VAF vs IS-determined LOB values**. For each alt displayed, the PET score from panel a was used to estimate the IS-determined Limit of Blank (LOB) (x-axis) which was plotted relative to VAF (y-axis); symbols as described for panel b. Dashed line is the 1:1 correlation in IS-derived LOB and corresponding NT VAF values.

### SNAQ-SEQ pipeline analysis improved clinical sensitivity without loss of specificity

In *Sample EfIS, Sample A* was 25-fold diluted with *Sample B* (5-fold greater dilution with *Sample B* than in *Sample DfIS*). Among the four replicate *Sample EfIS* libraries, 80 ground truth TP variant measurements were expected. However, because the mean TP VAF in *Sample EfIS* value was expected to be less than 0.5%^30^, this sample presented a stringent test for clinical sensitivity of NGS measurement platforms to measure the ground truth TP variants. The effect of SNAQ-SEQ QC with PET analysis on ILM UMI NGS analysis ground truth TP variants in *Sample EfIS* replicate libraries is presented in **Table 1**. Some TP variants were not detected by ILM (i.e. no alts with either VCF PASS or LowSupport FILTER) and are represented as blank cells. The TP that were called significant by SNAQ-SEQ QC with PET analysis but not called by ILM UMI NGS alone (i.e., FILTER LowSupport but not PASS) are indicated as VAF values with green background. One *Sample EfIS* variant called by ILM (i.e., VCF PASS FILTER) was not supported by SNAQ-SEQ QC analysis and is indicated as a VAF value with red background. The remaining cells contain VAF values of variants that were called by both methods (i.e., Illumina VCF PASS FILTER and significant by SNAQ-SEQ QC PET Analysis SNAQ-SEQ QC analysis of ILM VCF file variants with LowSupport FILTER annotation did not result in any false positive calls.

**Table 1.** Effect of SNAQ-SEQ QC on ILM UMI NGS Analysis of ground truth true positive variants in *Sample EfIS* replicate libraries. TP (True Positive) Variant Annotation: CHROM, POS, REF, ALT, GENE indicate hg19 chromosome location, reference base, alternate base and HUGO Gene Nomenclature Committee (HGNC) symbol; True Positive Variant allele fraction (VAF) (%) in Replicate (Rep) 1, 2, 3, 4; blank cells indicate variant not detected by Illumina pipeline (i.e. no alts with either VCF PASS or LowSupport FILTER), green-filled cells indicate alts not called by Illumina (i.e., VCF alts with LowSupport FILTER only) that were called by SNAQ-SEQ QC PET analysis, red-filled cell indicates an Illumina VCF alt called by Illumina (i.e., VCF PASS Filter) but not significant by SNAQ-SEQ QC PET analysis. Remaining variant cells contain variants that were called by both methods (i.e., Illumina VCF PASS FILTER and significant by SNAQ-SEQ QC PET Analysis).

| Table 1. Effect of SNAQ-SEQ QC on ILM UMI NGS Analysis of ground truth true positive variants in *Sample EfIS* replicate libraries |  |  |  |  |  |  |  |  |
| --- | --- | --- | --- | --- | --- | --- | --- | --- |
| True Positive Variant Annotation |  |  |  |  | True Positive Variant VAF |  |  |  |
| CHROM | POS | REF | ALT | GENE | Rep1 | Rep2 | Rep3 | Rep4 |
| chr1 | 115256529 | T | A | NRAS | 0.4% | 0.2% | 0.2% | 0.2% |
| chr1 | 115258748 | C | A | NRAS | 0.1% |  | 0.2% |  |
| chr12 | 112888162 | G | C | PTPN11 | 0.1% | 0.3% | 0.2% |  |
| chr15 | 66729162 | C | T | MAP2K1 |  | 0.1% | 0.4% | 0.2% |
| chr17 | 7576569 | G | A | TP53 |  | 0.3% | 0.2% | 0.2% |
| chr17 | 7577022 | G | A | TP53 | 0.3% | 0.2% | 0.2% | 0.1% |
| chr17 | 7577085 | C | T | TP53 | 0.1% | 0.2% | 0.1% | 0.1% |
| chr17 | 7577118 | C | A | TP53 | 0.5% | 0.5% | 0.3% | 0.5% |
| chr17 | 7577529 | A | T | TP53 | 0.6% | 0.4% | 0.5% | 0.2% |
| chr17 | 7578211 | C | T | TP53 | 0.4% | 0.3% | 0.4% | 0.4% |
| chr17 | 7578671 | C | T | TP53 | 0.2% | 0.2% | 0.5% |  |
| chr17 | 7578679 | A | G | TP53 | 0.3% | 0.5% | 0.7% | 0.5% |
| chr17 | 37880987 | C | T | ERBB2 |  | 0.2% | 0.2% | 0.2% |
| chr3 | 178936091 | G | A | PIK3CA | 0.2% | 0.2% | 0.3% | 0.1% |
| chr4 | 1803153 | G | A | FGFR3 | 0.2% | 0.2% | 0.2% | 0.2% |
| chr4 | 1803172 | G | A | FGFR3 | 0.2% | 0.2% | 0.2% | 0.2% |
| chr4 | 1803173 | C | T | FGFR3 | 0.3% | 0.1% | 0.1% | 0.2% |
| chr4 | 1803385 | G | C | FGFR3 | 0.2% | 0.2% | 0.2% | 0.3% |
| chr4 | 1803704 | T | C | FGFR3 | 0.8% | 0.7% | 0.6% | 0.6% |
| chr9 | 139399344 | A | G | NOTCH1 | 0.3% | 0.3% | 0.5% |  |

As indicated in the summary table (**Table 2**) the Illumina gVCF file used for SNAQ analysis <u>detected</u> all 80 TP in the four replicate libraries from *Sample EfIS*, but only <u>reported</u> (i.e. called) 64 as PASS (overall 80% sensitivity). The Illumina FILTER label for the remaining 16 TP was LowSupport (i.e., not reportable). In comparison, SNAQ-SEQ QC analysis called 71 of the 80 TP (89% sensitivity) for a 9% increase in sensitivity. Notably, all of the increase in sensitivity occurred among the very low VAF TP variants in *Sample EfIS* (0.1% ≤ VAF ≤ 0.3%). Among these TP variants the Illumina pipeline called 69% of TP in the ILM VCF file with PASS FILTER, whereas SNAQ-SEQ QC analysis of TP in the same ILM VCF file with either PASS or LowSupport FILTER increased TP detection sensitivity to 81%, for a 13% increase in sensitivity.

**Table 2.** Summary: SNAQ effect on detection sensitivity. Detection rate of ground truth TP variants by ILM variant caller (i.e., variants with ILM VCF PASS FILTER), or ILM caller plus SNAQ-SEQ QC (SNAQ) caller for variants with ILM VCF PASS + LowSupport FILTERS. ILM and SNAQ variant callers were compared among variants with low, high, or all mean inter-replicate VAF values: VAF ≥ 0.10% and ≤ 0.3%, > 0.3% VAF, and ALL VAF variants. Sensitivity was calculated by dividing the number of *Measured* variants by *Expected* variants, with the increase in sensitivity indicated in *SNAQ Sensitivity Increase* row.

| Table 2. Summary: SNAQ effect on detection sensitivity |  |  |  |  |  |  |
| --- | --- | --- | --- | --- | --- | --- |
| Variant VAF Range | $0.10\% \leq \text{VAF} \leq 0.3\%$ | | $\text{VAF} > 0.3\%$ | | All VAF | |
| Variant Caller | ILM | SNAQ | ILM | SNAQ | ILM | SNAQ |
| Expected TP calls | 56 | 56 | 24 | 24 | 80 | 80 |
| Observed TP calls | 41 | 48 | 23 | 23 | 64 | 71 |
| Sensitivity | 73% | 86% | 96% | 96% | 80% | 89% |
| SNAQ Sensitivity Increase | 13% |  | 0% |  | 9% |  |

## DISCUSSION

Inclusion of SNAQ-SEQ IS in each contrived ctDNA sample prior to library preparation enabled PET analysis and calculation of an LOB, LOD, and significance score for each variant at each targeted nucleotide position in each sample. SNAQ-SEQ QC was validated in NGS following either non-UMI or UMI library preparation and increased overall clinical sensitivity to call TP variants relative to Illumina UMI pipeline alone (**Tables 1, 2**). The TP variant clinical sensitivity benefit was greatest for *Sample EfIS*, in which the median VAF measured for TP variants was 0.25%. Notably, for TP variants with VAF ≤ 0.3% the sensitivity with ILM UMI pipeline alone was 69% and SNAQ-SEQ QC increased sensitivity to 82% with no decrease in clinical specificity (**Table 2)**. It should be noted that typical ctDNA clinical specimens contain about 25 ng (7,500 genome copies). Thus, depending on library capture efficiency for each target, 25 ng could support report of variants as low as 0.1% VAF using the three deduplicated read detection limit applied in this SNAQ-SEQ QC study. Further, efforts to validate below 0.1% VAF would not have practical meaning with respect to analysis of clinical ctDNA specimens.

Use of SNAQ-SEQ QC to report variants based on calculation of an LOB, LOD, and significance score for each variant at each targeted nucleotide position in each sample is a significant advance compared to current practice in which a predefined VAF LOD is imposed^11, 30^ or reports are restricted to measurements that reach a threshold coverage across the targeted regions. In the absence of SNAQ-SEQ QC, poor accuracy for calling variants with VAF <0.5% may be observed due to a) variable low yield from library preparation, b) variable loss of signal associated with stochastic sampling and c) low VAF signal to noise (i.e., position- and nucleotide-change specific NGS background technical error). Variant callers (e.g. VarDict) based on statistical analysis alone are tuned to a narrow range of coverage and assumption of NGS error rates based on panels of “normal” samples. In contrast, the SNAQ-SEQ QC significance scores based on PET analysis adjust to sample- and target-specific coverage and directly tested error rate. Therefore, this approach will support variant calling in sub-optimal sequence data resulting from low genomic input or low insert yields due to sample quality or unforeseen variation in reagents or equipment function.

In this and prior studies there was no evidence that inclusion of SNAQ-SEQ IS in samples interfered with testing sensitivity or specificity. The SNAQ-SEQ IS mixture used in this study was designed to span each known actionable mutation. This IS mixture spanned 4% of the TST170 panel, thus there was no significant impact on sample coverage. Further, mixture of SNAQ-SEQ IS with samples had no effect on workflow or performance of established targeted NGS platforms in an accompanying SEQC2 study^30^.

Taken together, these findings demonstrate that application of synthetic spike-in IS will provide critical test- and sample-specific quality control in NGS diagnosis of actionable mutations, as they do in other key molecular diagnostic testing methods, including liquid and gas chromatography, mass spectroscopy^32, 33^ and the FDA-approved Roche COBAS^®^ qPCR tests^34^.

This, study revealed analytical performance characteristics of the SNAQ-SEQ QC method that may be improved. First, there was systematic inter-platform variation in hybrid capture of IS relative to NT (data not shown). That said, once the appropriate IS input to achieve the desired IS to NT ratio (i.e. >2.5 IS:NT) was empirically identified for a hybrid capture platform, that platform yielded an inter-experimentally consistent result (**Supplemental Table 1**). Therefore, until the basis for this systematic bias is better understood, the protocol will be to experimentally determine for each hybrid capture panel platform the IS DNA input required to achieve the desired IS:NT ratio following library preparation. Second, higher IS coverage increases power to establish LOB based on PET, due to the larger number of <u>potential</u> observations for technical error. We empirically determined that an IS:NT ratio of at least 2.5:1 provided optimal power to measure LOB by PET across all positions/nucleotide exchanges represented in this study.

In summary, the SNAQ-SEQ orthogonal biochemical/statistical approach to NGS QC improved clinical sensitivity for measurement of ground truth positive variants in SEQC2 contrived ctDNA reference samples without loss of clinical specificity. This advance was possible due to increased analytical specificity. Specifically, use of SNAQ-SEQ QC enabled determination of technical error, LOB, confidence limits and calculation of lower LOD for each variant, at each nucleotide position, in each sample. This represents a major advance because, due to the limited number of genome copies in ctDNA specimens, it is critical to maximize the ability to reliably report low VAF actionable mutations. This approach controls for systematic chemical-physical NGS errors, pipeline-specific non-systematic errors, and coverage. Our work promises to provide a reliable method to increase the yield of reportable variant calls in low input clinical ctDNA samples.

## METHODS

The aim of this study was to a) design methods for incorporation of synthetic human genome reference sequence internal standard spike-ins into hybrid-capture targeted-NGS UMI library analysis, b) validate these methods as orthogonal QC in SEQC2 *Sample A* following Roche SeqCap EZ Choice custom PHC panel enrichment, positional deduplication, and Poisson Exact Test (PET) analysis of NT variant reads relative to IS variant reads, then c) evaluate clinical utility of this method in SEQC2 contrived ctDNA *Samples DfIS* and *EfIS* following Illumina TST170 hybrid capture panel enrichment, UMI deduplication, and PET analysis of NT variant and IS variant reads.

### Design and synthesis of reference DNA internal standard [IS] spike-ins for use in hybrid capture NGS

We prepared a mixture of synthetic Standardized Nucleic Acid Quantification for Sequencing (SNAQ-SEQ) IS DNA controls that span 38 kb and correspond to 32 exons and 61 actionable mutations^12, 35, 36^. This IS mixture was used successfully in previous work to provide quality control in amplicon libraries prepared with DNA extracted from tissue samples^12^. To provide quality-control for targeted hybrid capture library analysis of SEQC2 contrived ctDNA reference materials used in this study, the IS mixture was enzymatically sheared to approximate the modal distribution of typical ctDNA samples.^30^ Of this 38kb region, 6.8 kb overlap with the Roche SeqCap EZ Choice custom PHC panel used to establish ground truth in *Sample A* and the Illumina TST170 hybrid capture panel used to analyze the SEQC2 contrived ctDNA *Samples DfIS* and *EfIS*.

Each IS sequence matched the respective hg19 human reference genome sequence except for unique dinucleotides (DN) every 40-60 bp to allow for bioinformatic separation of IS from native template (NT) genomic reads. Each IS control spanned sequence that comprised one or more full exons containing the actionable mutation(s) plus up to 500 bp of DNA flanking each side of each mutation. Flanking DNA was included in an effort to ensure that enzymatic or mechanical fragmentation methods would yield a size spectrum around each targeted mutation that closely approximated that of sample DNA. The DN positions were chosen to avoid actionable mutation sites, known primer binding sites used by commercial PCR amplicon library vendors, and known genomic variants with VAF ≥ 0.1% based on HapMap database^37^. Each IS control was cloned into a pUC plasmid and the sequence was verified. Other than the DN sites, it was expected that every IS position matched reference genome due to the low replication error rate of *E. coli* DNA, which is <1 x 10^−8^. Thus, spanning 38,000 bp the expected error rate would be < 0.004%. Plasmids were linearized, quantified and mixed at an equimolar concentration, with <20% CV for inter-IS variation in mixture. For this study, a large batch of IS mixture was then enzymatically fragmented with goal to achieve a 130-170 bp modal size distribution, closely approximating that observed in typical ctDNA clinical specimens as well as the contrived SEQC2 ctDNA reference materials^30^. Frozen aliquots of IS mixture were then distributed to participating SEQC2 sites for use in this and related SEQC2 studies^30^.

### SEQC2 Reference Samples

Reference *Samples A* and *B* used in this study were developed by the SEQC2 Oncopanel Sequencing Working Group and described in related studies from the SEQC2 project^31^. SEQC2 *Sample A* comprised a mixture of ten cancer cell lines, with an expected VAF for most variants ranging from 2% to 15%. Lower VAF samples were created by mixing *Samples A* and *B*. Briefly, *Sample A* was combined with DNA extracted from a non-cancer background cell-line (*Sample B*) to create two additional reference samples, *Sample D* (1:5 dilution; 20% *Sample A* / 80% *Sample B*), and *E* (1:25 dilution; 4% *Sample A* / 96% *Sample B). Samples B, D*, and *E* then were enzymatically fragmented to create *Samples Bf, Df*, and *Ef* with modal 130-170 bp fragment size^30^. *Samples Bf, Df*, and *Ef* were mixed with the fragmented IS spike-in mixture to make SEQC2 test *Samples BfIS, DfIS* and *EfIS*, respectively. Each of these test samples were then distributed to each test site that participated in this study. With this design it was intended that following mixture with a sample, each IS DNA sequence would closely parallel its corresponding sample sequence during hybrid capture library preparation and sequencing yet be easily separated bioinformatically from sample sequence during pipeline analysis.

### Establishment of ground truth dataset

*Samples A* and *B* had limited ground truth data for the TST170 x SNAQ-SEQ IS overlap available from the SEQC studies^31^. Because this would limit the assessment of SNAQ-SEQ controls as an independent NGS quality control, a more complete ground truth was created. Specifically, SEQC2 *Sample A* was serially two-fold diluted with *Sample B* (normal subject DNA)^31^ to create 1, 2, 4, 8, 16, 32 and 64-fold dilutions. Then, each serial dilution sample was mixed with SNAQ-SEQ synthetic spike-in controls to achieve IS:NT ratio of >2.5 for each target^31^. Following enzymatic fragmentation, triplicate aliquots of each sample were enriched on the Roche SeqCap EZ Choice custom PHC panel followed by non-UMI pipeline library preparation then sequenced on an Illumina Novaseq S4 at the University of Texas Southwestern. As schematically depicted in **Figure 1a**, the FASTQ files were shipped on hard drives to AccuGenomics, Inc., reads split using custom PERL script described above, aligned with BWA mem 0.7.17-r1194-dirty, deduplicated with GATK markDuplicates v4.0.11.0 and a genomic (g)VCF file generated with VarDict 1.5.7 for each sample. Ground truth positive (TP) variants in *Sample A* were defined as those for which VAF responded to serial dilution. The serial dilution VAF results are presented in **Supplemental Figure 1**. The positionally deduplicated coverage for each target was above 1,100 for each replicate, and this coverage was sufficient to confirm that a position was ground truth negative (TN) down to at least 0.5% VAF. The goal was to both establish ground truth positive (TP) variants and confirm that any other variant was a false positive (FP) by confirming lack of titration response. Data from this study were used to develop the SEQC-SEQ QC method, as presented in **Figure 1c-d** and presented in **Results** section.

### Illumina TST170 hybrid capture NGS library preparations and analysis

50 ng aliquots of *Samples BfIS, DfIS* and *EfIS* were each included in four replicate TST170 UMI libraries prepared at Q2 Solutions, Inc. according to the ILM2 method previously described ^30^, and sequenced on NextSeq. Following target enrichment, library preparation, and sequencing, NT and IS reads from *Samples BfIS, DfIS*, or *EfIS*, were bioinformatically isolated into their respective bins. Specifically, resulting FASTQ files were separated into NT, IS and suspicious using a custom designed SIST3.1.4.pl PERL script prior to running each sequence pool into the analysis pipeline (**Figure 1a**). Suspicious reads had both NT and IS base changes in the same read, made up 0.1% of reads, and likely arose by sequencing error or recombination between NT and IS during library preparation.

### Illumina UMI Pipeline and Variant Calling

After bioinformatic isolation of NT and IS reads into different FASTQ files at AccuGenomics the NT and IS FASTQ files were uploaded to BaseSpace^38^. Each NT FASTQ file was processed through the standard protocol which yielded a UMI deduplicated BAM file and VCF file. Each IS FASTQ file was processed through the same protocol except for alignment to the hg19 human reference genome modified to include the IS-specific dinucleotide changes, which yielded UMI deduplicated BAM files and VCF files. Because each IS was known to be reference sequence, any variant alleles were, by definition, technical error. Analysis of the IS VCF file indicated certain artifactual variants that were not detected by the Illumina variant caller. These variants likely were created during fragment end repair from 3-prime self-priming events and it was determined empirically that a VarDict PMEAN filter effectively removed them. Specifically, the VarDict variant caller PMEAN score indicated the average distance a variant occurred from a fragment end. To eliminate these end artifacts, VarDict analysis generated a PMEAN score for each variant in UMI deduplicated BAM files from each of 12 libraries (four replicate libraries each for *Samples BfIS, DfIS* and *EfIS*), then an R-script was used to calculate the median PMEAN score for each variant across all 12 samples. Analysis of the Illumina VCF file variants indicated that a median PMEAN of 25 would separate the fragment end repair artifact variants from TP variants (**Supplemental Figure 3**). The <25 PMEAN positions were blacklisted from further analysis, reducing the total IS and NT variant sequences by 10%. Following application of the PMEAN 25 filter, Illumina UMI deduplicated BAM files were used to generate VCF files that reported all variants and variant positions. Within each Illumina VCF file, called variants were annotated as FILTER PASS, and statistically less likely non-called variants were annotated with FILTER LowSupport, Blacklist, or LowVar.

### SNAQ-SEQ QC variant calling of Illumina UMI deduplicated BAM files

At AccuGenomics, SNAQ-SEQ QC analysis was combined with Poisson Exact Test (PET) to calculate the statistical difference between each sample library NT variant VAF and respective IS variant VAF based on NT variant count and position coverage and IS count and position coverage. SNAQ-SEQ QC with PET analysis was performed on any variant meeting the following criteria: a) not in Illumina’s blacklist, b) not in the PMEAN blacklist, c) ≥ three deduplicated reads, and 4) Illumina VCF FILTER containing PASS or LowSupport.

Because the IS was reference sequence prior to library preparation, it was assumed that any IS variant observed following library preparation and sequencing resulted from technical error. The 5% alpha used as the cutoff for NT variant VAF significance was adjusted using a Bonferroni correction, 0.05 divided by the number of the sample NT variants being tested (N=20 for each replicate library of *Sample EfIS*). An NT variant with VAF PET score less than the cutoff indicated the genomic variant passed SNAQ-SEQ QC and was called. An NT variant VAF PET score greater than the significance cutoff was not distinguishable from NGS error and was not called. SNAQ-SEQ analysis of IS, which treats the IS as the “sample” and the NT sequence as “background” was used to examine the test pipeline performance, as a working pipeline should yield no statistically significant IS variants.

### Performance metrics

Traditionally, a false negative (FN) is a ground truth positive (TP) variant not detected by sequence analysis, and a false positive (FP) is a variant detected at a ground truth negative (TN) site by sequence analysis. With SNAQ-SEQ QC, a FN could arise when a TP variant did not have a significant PET score, and a FP may arise if the observed NT variant derives from technical error in the sample that does not covary to the same extent in the IS.

## ACKNOWLEDGEMENTS

All SEQC2 participants freely donated their time, samples, reagents, and computing resources for the completion and analysis of this project. We thank SEQC2 Oncopanel Sequencing Working Group participants for providing useful feedback during manuscript preparation. This work was supported by grants from the National Cancer Institute (5U24CA086368; 1U01CA243483), the FDA (BAA grant HHSF223201510172C), the National Health and Medical Research Council (NHMRC) of Australia (grants APP1108254, APP1114016, APP1173594, and Cancer Institute NSW Early Career Fellowship 2018/ECF013). None of these funding bodies had a role in the design of the study, the collection, analysis, and interpretation of data, or in writing of the manuscript.

## AUTHORS’ CONTRIBUTIONS

JCW contributed to conception and design of the study, analysis and interpretation of data, and drafting and revision of the manuscript.

TM contributed to conception and design of the study, acquisition, analysis and interpretation of data; the creation of new software used in the work and drafting and revision of the manuscript.

BA contributed to acquisition and analysis of the data and the creation of new software used in the work.

ELC contributed to conception and design of the study, acquisition and interpretation of data, and revision of the manuscript.

DJC contributed to conception and design of the study, acquisition, analysis and interpretation of data, the creation of new software used in the work and revision of the manuscript.

TMB contributed to conception and design of the study, the creation of new software used in the work. and revision of the manuscript.

WDJ contributed to acquisition of data and revision of manuscript.

AW contributed to acquisition of data.

JSL contributed to acquisition, analysis, and interpretation of data and revision of the manuscript

NH contributed to acquisition, analysis, and interpretation of data.

TAR contributed to acquisition, analysis, and interpretation of data and revision of the manuscript

NN contributed to acquisition, analysis, and interpretation of data and revision of the manuscript.

RK contributed to interpretation of data and revision of the manuscript.

GC contributed to acquisition, analysis, and interpretation of data and revision of the manuscript.

QZL contributed to acquisition, analysis, and interpretation of data and revision of the manuscript.

DJ contributed to analysis and interpretation of data and revision of the manuscript.

IWD contributed to interpretation of data and revision of the manuscript.

TM contributed to interpretation of data and revision of the manuscript.

LW analysis and interpretation of data, the creation of new software used in the work and revision of the manuscript.

JX contributed to conception and design of the study, analysis and interpretation of data, the creation of new software used in the work and revision of the manuscript.

## SUPPLEMENTAL INFORMATION

**Supplemental Figure 1.**
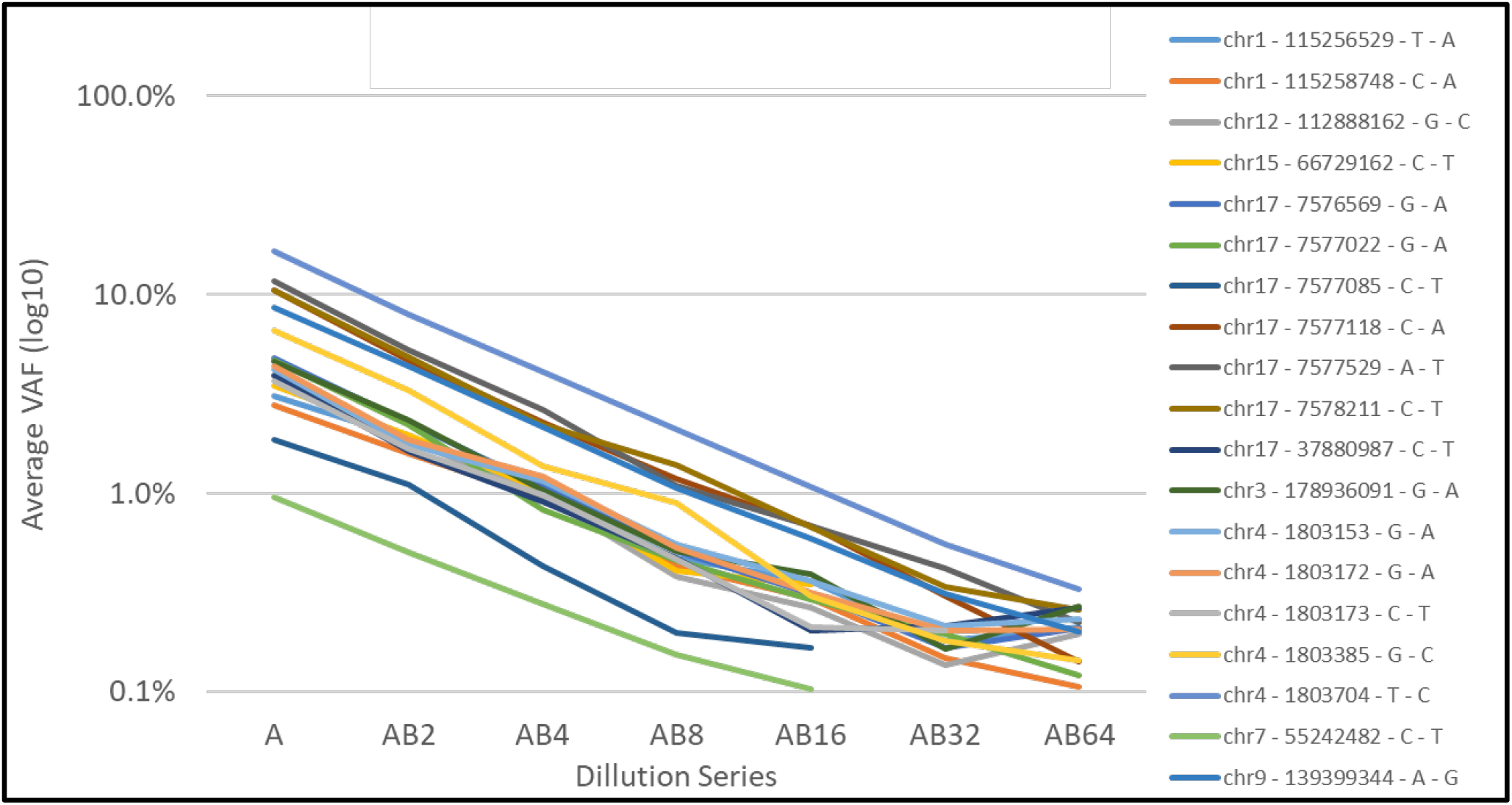
Identification of *Sample A* ground truth true positive (TP) Variants. Ground truth TP variants were identified within the consensus region for the Roche SeqCap EZ Choice custom PHC panel TST170 panel and SNAQ-SEQ IS panel, based on serial dilution of *Sample A* with *Sample B* from 2- to 64-fold (AB2 through AB64). Plotted are the average replicate VAF values for 19 variants that demonstrated a dilution response, with information for each variant in the legend. One additional *Sample A* TP variant restricted to the consensus region between TST170 and SNAQ-SEQ IS (i.e., not in Roche panel) was identified through serial dilution (data not shown), resulting in a total of 20 ground truth TP variants. (see **Table 2**).

**Supplemental Figure 2.**
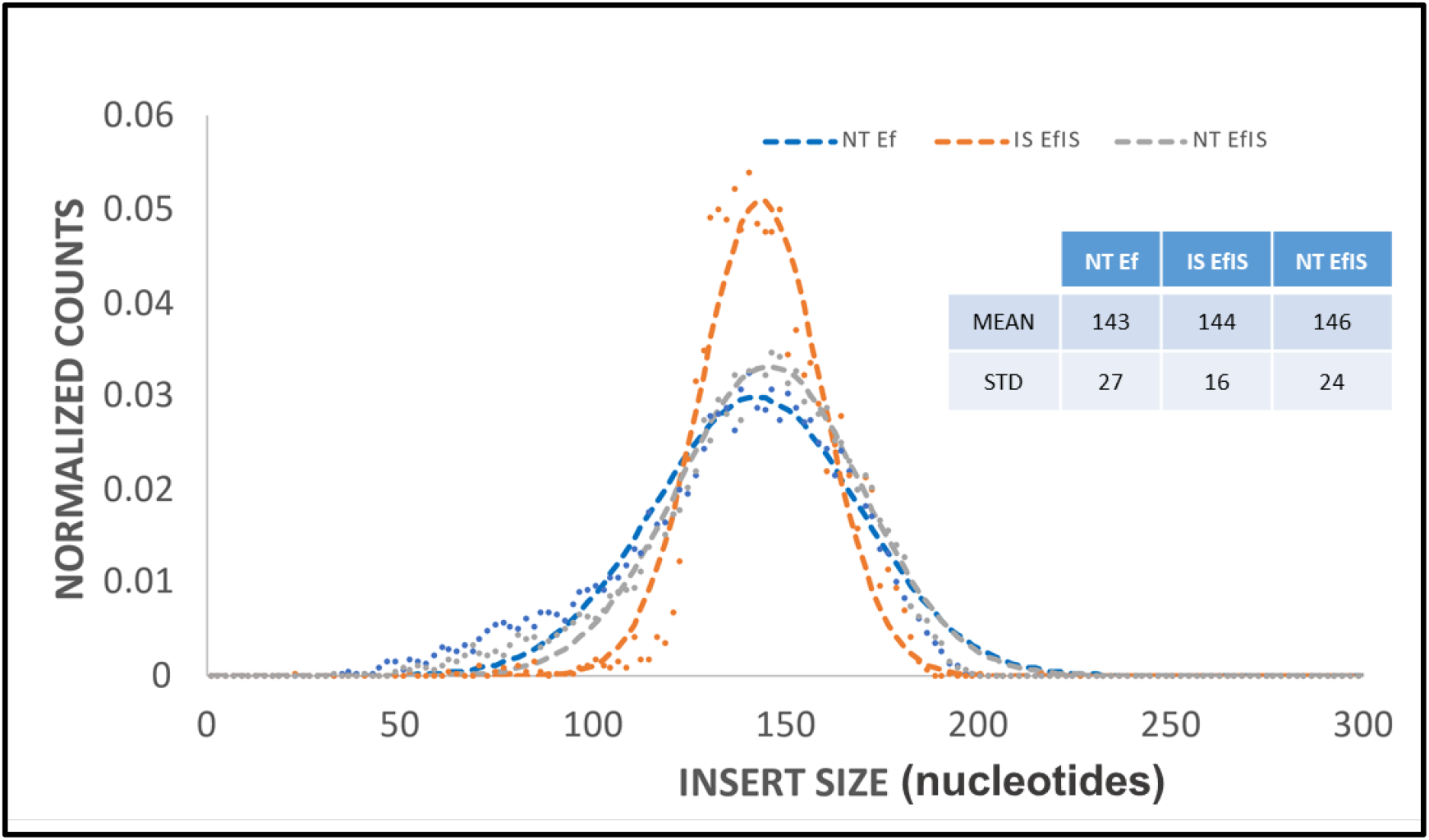
*Samples Ef* and *EfIS* sequence read size distribution for native template (NT) and (IS). Sequencing libraries were prepared for *Samples Ef* and *EfIS*. Following sequencing and bioinformatic separation of NT and IS sequence reads derived from the *Sample EfIS* library, the sequence read size distributions were plotted as histograms. x-axis: sequence insert size in nucleotides; y-axis: normalized counts as a fraction of total. Table insert: Mean and standard deviation (STD) sequence read length values for NT in Sample *Ef*, IS in *Sample EfIS*, and NT in *Sample EfIS*.

**Supplemental Figure 3.**
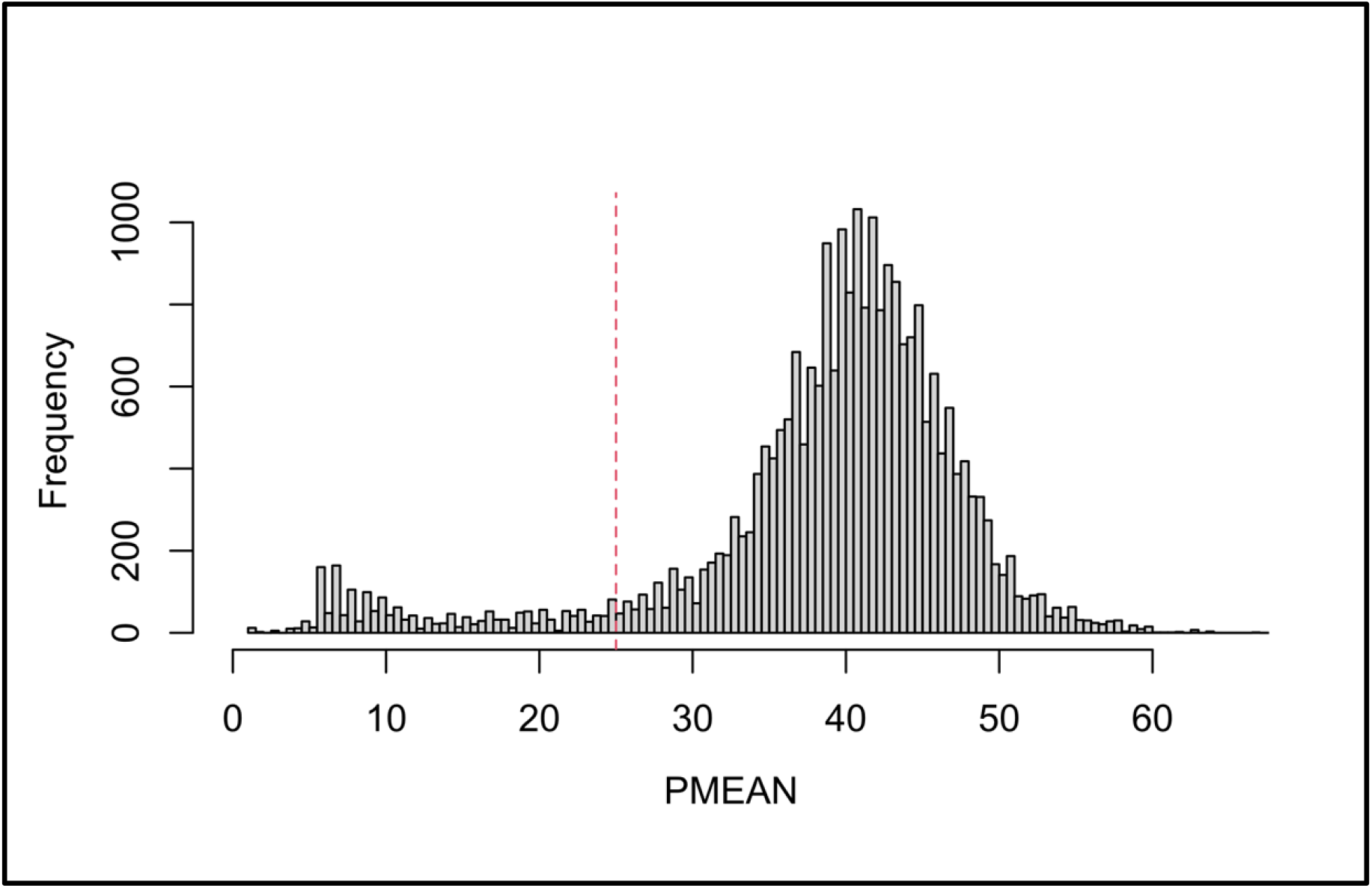
Application of PMEAN filter to remove non-systematic errors near fragment ends that are not suppressed by UMI. Displayed are all ILM VCF alts with FILTER containing PASS or LowSupport among UMI deduplicated BAM files for each replicate library for *Samples BfIS, DfIS*, and *EfIS* (12 libraries). PMEAN (x-axis): An R-script was used to determine aggregated median PMEAN value for indicated alt bins. Frequency (y-axis): the number of alts in each bin. The red dashed horizontal line indicates the PMEAN 25 threshold used to detect end repair artifacts with alts in positions below this threshold removed from analysis.

**Supplemental Table 1.** Average IS/NT ratio for each IS in the SNAQ-SEQ IS mixture. The IS mixture comprised 32 IS derived from 27 plasmids with plasmid IDs indicated. Each plasmid contained IS for one or more exons within a gene. For each IS plasmid, the IS:NT ratio mean, SD, and CV was derived from the IS:NT ratio across each position shared by each SNAQ-SEQ IS in the plasmid and the TST170 panel, for each of 12 libraries including four replicates of three samples: *Samples BfIS, DfIS*, and *EfIS*.

| Supplemental Table 1. Average IS/NT ratio for each IS in the SNAQ-SEQ IS mixture |  |  |  |
| --- | --- | --- | --- |
| IS Plasmid ID | IS:NT Ratio |  |  |
|  | Mean | SD | CV |
| JAK3 | 2.9 | 0.14 | 5% |
| MAP2K1_3 | 3.4 | 0.50 | 14% |
| NRAS_3 | 3.6 | 0.18 | 5% |
| PDGFRA_12 | 3.7 | 0.11 | 3% |
| ERBB2_RST | 3.8 | 0.18 | 5% |
| NRAS_2 | 3.9 | 0.35 | 9% |
| EGFR19_RST | 4.0 | 0.13 | 3% |
| PTEN | 4.2 | 0.30 | 7% |
| EGFR21_RST | 4.4 | 0.25 | 6% |
| KRAS_3 | 4.4 | 0.17 | 4% |
| JAK2_14 | 4.5 | 0.31 | 7% |
| ATM | 4.5 | 0.21 | 5% |
| FGFR2_7 | 4.6 | 0.12 | 3% |
| EGFR18_RST | 4.7 | 0.32 | 7% |
| PIK3CA_10 | 4.8 | 1.00 | 21% |
| JAK2_12 | 4.8 | 0.67 | 14% |
| EGFR20_RST | 4.8 | 0.26 | 5% |
| NOTCH1 | 5.0 | 0.76 | 15% |
| TP53 | 5.1 | 0.74 | 14% |
| MET | 5.2 | 0.37 | 7% |
| PTPN11 | 5.6 | 0.12 | 2% |
| FBXW7_16 | 6.0 | 0.18 | 3% |
| CTNNB1 | 6.0 | 0.47 | 8% |
| FGFR3 | 6.1 | 0.58 | 9% |
| KIT_17 | 6.1 | 0.71 | 11% |
| KIT_9 | 6.8 | 0.54 | 8% |
| FBXW7_13 | 7.3 | 0.28 | 4% |
| All Plasmid MEAN | 4.8 | 0.37 | 7.6% |

## References

1. Beaubier, N. et al. Clinical validation of the tempus xT next-generation targeted oncology sequencing assay. Oncotarget 10, 2384–2396 (2019).

2. Chen, H.Z., Bonneville, R. & Roychowdhury, S. Implementing precision cancer medicine in the genomic era. Semin Cancer Biol 55, 16–27 (2019).

3. Schwaederle, M. et al. Association of Biomarker-Based Treatment Strategies With Response Rates and Progression-Free Survival in Refractory Malignant Neoplasms: A Meta-analysis. JAMA Oncol 2, 1452–1459 (2016).

4. Rodon, J. et al. Genomic and transcriptomic profiling expands precision cancer medicine: the WINTHER trial. Nat Med 25, 751–758 (2019).

5. Zutter, M.M. et al. The Cancer Genomics Resource List 2014. Arch Pathol Lab Med 139, 989–1008 (2015).

6. Siravegna, G., Marsoni, S., Siena, S. & Bardelli, A. Integrating liquid biopsies into the management of cancer. Nat Rev Clin Oncol 14, 531–548 (2017).

7. Vasan, N., Baselga, J. & Hyman, D.M. A view on drug resistance in cancer. Nature 575, 299–309 (2019).

8. Volckmar, A.L. et al. A field guide for cancer diagnostics using cell-free DNA: From principles to practice and clinical applications. Genes Chromosomes Cancer 57, 123–139 (2018).

9. Corcoran, R.B. & Chabner, B.A. Application of Cell-free DNA Analysis to Cancer Treatment. The New England journal of medicine 379, 1754–1765 (2018).

10. Oxnard, G.R. et al. Association Between Plasma Genotyping and Outcomes of Treatment With Osimertinib (AZD9291) in Advanced Non-Small-Cell Lung Cancer. Journal of clinical oncology: official journal of the American Society of Clinical Oncology 34, 3375–3382 (2016).

11. https://www.accessdata.fda.gov/cdrh_docs/pdf20/P200010A.pdf (2020).

12. Craig, D.J. et al. Technical advance in targeted NGS analysis enables identification of lung cancer risk-associated low frequency TP53, PIK3CA, and BRAF mutations in airway epithelial cells. BMC cancer 19, 1081 (2019).

13. Blomquist, T., Crawford, E.L., Yeo, J., Zhang, X. & Willey, J.C. Control for stochastic sampling variation and qualitative sequencing error in next generation sequencing. Biomol Detect Quantif 5 (2015).

14. Fu, G.K. et al. Molecular indexing enables quantitative targeted RNA sequencing and reveals poor efficiencies in standard library preparations. Proc Natl Acad Sci 111 (2014).

15. Merker, J.D. et al. Circulating Tumor DNA Analysis in Patients With Cancer: American Society of Clinical Oncology and College of American Pathologists Joint Review. Journal of clinical oncology: official journal of the American Society of Clinical Oncology 36, 1631–1641 (2018).

16. Kuderer, N.M., Burton, K.A., Blau, S. & et al. Comparison of 2 commercially available next-generation sequencing platforms in oncology. JAMA Oncology (2016).

17. Stetson, D. et al. Orthogonal Comparison of Four Plasma NGS Tests With Tumor Suggests Technical Factors are a Major Source of Assay Discordance. JCO Precision Oncology, 1–9 (2019).

18. Rossi, G. & Ignatiadis, M. Promises and Pitfalls of Using Liquid Biopsy for Precision Medicine. Cancer research 79, 2798–2804 (2019).

19. Torga, G. & Pienta, K.J. Patient-Paired Sample Congruence Between 2 Commercial Liquid Biopsy Tests. JAMA Oncol 4, 868–870 (2018).

20. Squillace, R.M., Frampton, G.M., Stephens, P.J., Ross, J.S. & Miller, V.A. Comparing two assays for clinical genomic profiling: the devil is in the data. OncoTargets and therapy 8, 2237–2242 (2015).

21. Gargis, A.S. et al. Good laboratory practice for clinical next-generation sequencing informatics pipelines. Nature biotechnology 33, 689 (2015).

22. Blomquist, T.M. et al. Targeted RNA-sequencing with competitive multiplex-PCR amplicon libraries. PloS one 8, e79120 (2013).

23. https://www.fda.gov/science-research/bioinformatics-tools/microarraysequencing-quality-control-maqcseqc#MAQC_IV (2019).

24. Newman, A.M. et al. An ultrasensitive method for quantitating circulating tumor DNA with broad patient coverage. Nat Med 20, 548–554 (2014).

25. Cibulskis, K. et al. Sensitive detection of somatic point mutations in impure and heterogeneous cancer samples. Nature biotechnology 31, 213–219 (2013).

26. Kennedy, S.R., Salk, J.J., Schmitt, M.W. & Loeb, L.A. Ultra-sensitive sequencing reveals an age-related increase in somatic mitochondrial mutations that are inconsistent with oxidative damage. PLoS genetics 9, e1003794 (2013).

27. Schmitt, M.W. et al. Detection of ultra-rare mutations by next-generation sequencing. Proceedings of the National Academy of Sciences of the United States of America 109, 14508–14513 (2012).

28. Newman, A.M. et al. Integrated digital error suppression for improved detection of circulating tumor DNA. Nature biotechnology 34, 547–555 (2016).

29. Sandmann, S. et al. Evaluating Variant Calling Tools for Non-Matched Next-Generation Sequencing Data. Scientific reports 7, 43169 (2017).

30. Deveson, I. Evaluating the analytical validity of circulating tumor DNA sequencing assays for precision oncology. Nature biotechnology (in press).

31. Jones, W.D. A Verified Genomic Reference Material for Assessing Performance of Cancer Panels Detecting Small Variants of Low Allele Frequency. Genome Biol (in press).

32. Dolan, J.W. When should an internal standard be used? LCGC North America 30, 316–322 (2012).

33. Takats, Z., Wiseman, J.M. & Cooks, R.G. Ambient mass spectrometry using desorption electrospray ionization (DESI): instrumentation, mechanisms and applications in forensics, chemistry, and biology. J Mass Spectrom 40, 1261–1275 (2005).

34. Diagnostics, R. Roche Molecular Systems: cobas EGFR Mutation Test v2 https://diagnostics.roche.com/us/en/products/params/cobas-egfr-mutation-test-v2.html. (2015).

35. https://assets.thermofisher.com/TFS-Assets/LSG/manuals/MAN0017064_PanCancer_UG.pdf (2020).

36. Liu, J. et al. An Integrated TCGA Pan-Cancer Clinical Data Resource to Drive High-Quality Survival Outcome Analytics. Cell 173, 400–416 e411 (2018).

37. Sherry, S.T., Ward, M. & Sirotkin, K. dbSNP-database for single nucleotide polymorphisms and other classes of minor genetic variation. Genome research 9, 677–679 (1999).

38. https://www.illumina.com/products/by-type/informatics-products/basespace-sequence-hub.html, I.B. (2020).

